# An Integrated and Configurable End-to-End Pipeline for Longitudinal Cell Painting Analysis

**DOI:** 10.64898/2026.02.21.707179

**Authors:** Guang Zhao, Yu Xi, Mikhail Titov, Rebecca Weinberg, Sara Forrester, Tom Brettin, Shinjae Yoo, Xiaoning Qian, Byung-Jun Yoon

## Abstract

Cell painting assays generate high-dimensional, multi-channel imaging data that enable systematic characterization of cellular phenotypes. Increasingly, such assays are performed in longitudinal settings and under chronic perturbations, introducing additional challenges related to imaging variability, focus-field heterogeneity, and consistency across time points. Existing analysis workflows often require substantial manual adaptation to handle these complexities, limiting scalability and reproducibility. In this paper, we propose SCALE (Stable Cell painting Analysis for Longitudinal Experiments), an integrated, end-to-end analysis pipeline designed for robust longitudinal analysis of cell painting data. The pipeline combines nucleus-centered segmentation, automated quality control, feature extraction, and signal aggregation within a modular and configurable framework. Once assay-specific configurations are specified, the pipeline executes in a fully automated manner from raw images to downstream summary statistics and analysis-ready outputs. We demonstrate the utility of the pipeline using a chronic radiation exposure cell painting dataset, illustrating its ability to support consistent longitudinal comparisons across conditions and time points.

## 1 Introduction

Cell Painting assays combine multiple fluorescent stains with high-throughput microscopy to enable scalable, quantitative cellular phenotyping [3]. The resulting multi-channel images capture complementary aspects of cellular state across compartments, producing rich datasets that reflect morphological and organizational variation [4, 6]. The value of Cell Painting is amplified in large-scale and comparative experimental designs, where the goal is not to interpret a single image in isolation but to quantify reproducible differences across conditions [2, 12]. In these settings, downstream analysis depends critically on consistent processing segmentation, feature computation, aggregation, and summarization as small pipeline inconsistencies can obscure or distort the comparative signal.

Increasingly, Cell Painting experiments are designed as longitudinal studies or chronic perturbation assays, where the same system is imaged repeatedly over weeks and compared across multiple conditions. While standard Cell Painting workflows are often adequate for single time-point analysis, longitudinal settings introduce failure modes that are comparatively minor in one-off experiments but become dominant sources of fragility over time. First, longitudinal imaging commonly exhibits focus and z-stack variability across weeks, making intensity measurements sensitive to focal-plane selection and necessitating explicit rules for z-stack integration [13, 8]. Second, longitudinal acquisition comes with potential plate/batch drift and illumination shifts, requiring analysis outputs that remain comparable under baseline changes [13, 1]. Thirdly, many studies include heterogeneous staining panels within the same experiment, where subsets of wells use modified markers or channel substitutions, creating non-uniform channel semantics that must be handled explicitly [8, 7].

A rich ecosystem of Cell Painting software supports individual stages of analysis, including segmentation, feature extraction, and profile construction. For example, CellProfiler provides a modular image-analysis environment for segmentation and hand-crafted feature extraction [5, 14], DeepProfiler provides learned representation models for Cell Painting images [10], and pycytominer supports downstream profile processing (e.g., aggregation, normalization, and feature selection) from single-cell measurements [15]. However, in longitudinal experiments these workflows often become fragile because robustness must be maintained end-to-end, not just within a single module. First, common toolchains frequently require manual adaptation when imaging conditions, focus behavior, or staining panels vary across plates or weeks, which makes analyses difficult to scale and increases the risk of inadvertent bias. Second, many workflows do not enforce uniform execution across time points, allowing segmentation, filtering, or aggregation choices to adapt over the course of an experiment and undermining longitudinal comparability. Third, reproducibility is limited when key decisions are encoded in scripts or graphic user interface (GUI) settings rather than exposed through a configuration-driven interface that can be audited, versioned, and rerun deterministically. Finally, longitudinal analysis requires treating feature families differently: intensity readouts are particularly sensitive to focus and illumination and therefore need explicit integration rules, whereas structural descriptors (morphology/texture) have different stability properties. Existing workflows often do not make these feature-family differences first-class in the aggregation and reporting logic, leading to inconsistent handling of intensity versus structure over time.

In this paper we present SCALE (Stable Cell painting Analysis for Longitudinal Experiments) a stability-oriented, end-to-end pipeline for longitudinal Cell Painting analysis, focusing on integration, consistency and reproducibility. Rather than optimizing individual modules in isolation, we design the pipeline to operate as a unified system in which segmentation, quality control, feature extraction, aggregation, and reporting are explicitly coordinated. The workflow enforces condition-agnostic processing rules across time points and experimental conditions, reducing the risk of treatment-dependent analysis bias. Specifically, our contributions include:

- We present an integrated end-to-end pipeline that connects raw multi-channel images to longitudinal summary outputs, spanning segmentation, feature extraction, aggregation, and standardized reporting.
- We introduce a configuration scheme that supports longitudinal experimental structure and heterogeneous assay layouts.
- We develop a robustness-oriented segmentation module with automated QC and an adaptive, dataset-wide corrective loop.
- We demonstrate the pipeline on a chronic radiation Cell Painting assay spanning multiple dose levels and nine weeks of imaging to illustrate handling of heterogeneous staining panels, and longitudinal feature trajectories.

The remainder of this paper is organized as follows. Section 2 describes the integrated pipeline architecture; Section 3 demonstrates instantiation of the framework on a chronic radiation Cell Painting dataset; Section 4 concludes with a discussion of stability considerations in longitudinal imaging studies and outlines potential extensions.

## 2 The SCALE Framework

### 2.1 Pipeline Design Principles

The SCALE framework is designed to support robust, end-to-end analysis of longitudinal cell painting data acquired across multiple channels, focus fields, and experimental conditions. The pipeline emphasizes consistency among time points, modularity and configurable, enabling reuse across diverse assay designs. Overall, the pipeline includes configuration, segmentation, feature extraction and Result summary. The pipeline supports configuration setting, modular setting to combine different tools. The overall diagram of the pipeline is shown in Fig. 1.

**Figure 1.**
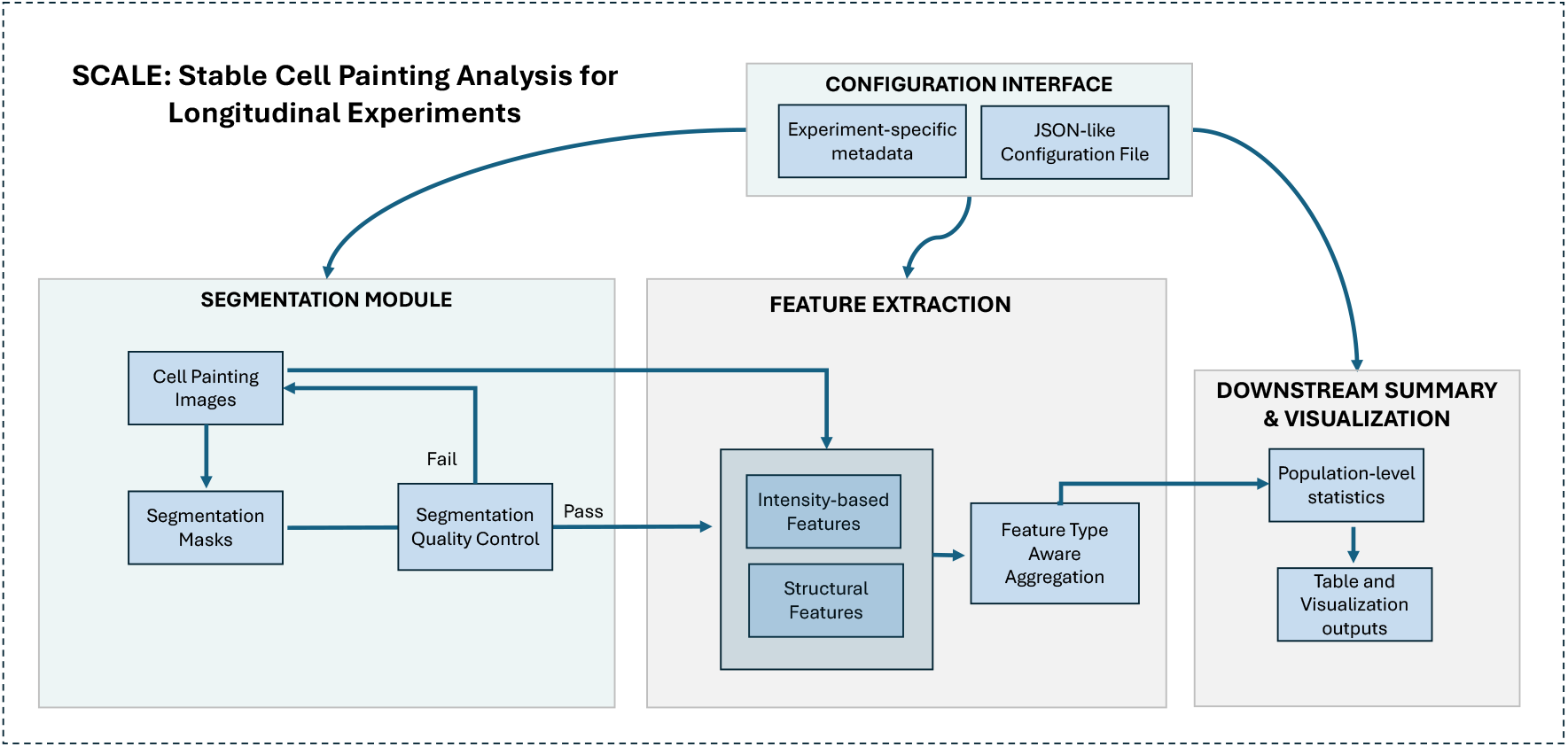
Overview of the SCALE framework for longitudinal Cell Painting analysis. The framework supports robust processing across multiple imaging channels, focus fields, time points, and experimental conditions. The pipeline is organized into four modular components: (1) a configuration interface that manages experiment-specific metadata and JSON-like configuration files; (2) a segmentation module that generates nucleus masks with quality control; (3) a feature extraction module that computes intensity-based and structural features followed by feature-type-aware aggregation; and (4) downstream summary and visualization, producing population-level statistics and tabulated outputs.

### 2.2 Configuration Interface

Cell painting assays often involve complex imaging configurations, include multiple fluorescence channels and focus fields per well. In longitudianl experiments, these configurations keeps consistent across time points, while the same analysis workflow is applied uniformly across all experimental conditions. In our pipeline, the assay-specific information is explicitly provided through a structured, JSON-like configuration interface. This configuration specifies what information is present in the dataset and how it should be interpreted by the pipeline, while the underlying processing logic remains unchanged. Such a design enables parallel execution of the pipeline under different configuration settings, facilitating systematic comparison of analysis choices.

Configurable elements include high-level categories such as the set of fluorescence channels available, the assignment of channels to intensity-based or structural feature extraction, the choice of segmentation backend, the selection of feature families to include, and the specification of downstream analysis outputs and visualization targets. Experiment-specific metadata, such as condition labels and time-point identifiers, are treated generically as categorical annotations, and they are used solely to define grouping keys for aggregation and comparison.

With this configuration setting, the proposed framework supports reuse across diverse cell painting experiments without requiring modification of the core analysis workflow.

### 2.3 Segmentation Module

Within SCALE, the segmentation module defines the cellular instances used throughout the pipeline and therefore serves as a foundational component for all downstream analyses. The primary objective of this module is robustness and consistency across time points, experimental conditions, and imaging settings, rather than maximizing per-cell accuracy under a single condition. Therefore, segmentation is implemented in a condition-agnostic manner and applied uniformly across the entire dataset.

#### 2.3.1 Segmentation Workflow

Segmentation is performed using a nuclear-guided strategy, in which nuclei are first identified from the nuclear staining channel. Nuclear signals typically exhibit high signal-to-noise ratios and stable spatial localization across imaging conditions, making them a reliable anchor for defining individual cellular instances.

The resulting nuclear masks serve as the primary reference for segmentation within the pipeline. For nuclear-localized feature extraction, these masks are used directly. For channels containing cytoplasmic or organelle-associated signals, cellular regions are derived in a secondary step based on the nuclear masks, for example by constrained expansion or region assignment methods. This nuclear-guided cellular segmentation provides approximate whole-cell regions while preserving consistent cell identity across channels. This approach is commonly adopted in cell painting and high-content imaging pipelines, where nuclear segmentation provides stable instance separation and supports downstream cellular feature extraction [5].

By prioritizing nuclear segmentation and treating cellular regions as derived representations, the pipeline avoids ambiguity in cell boundary definition that can arise from heterogeneous cytoplasmic signals, variable staining, or focus-dependent artifacts. All segmentation steps are executed uniformly across time points, experimental conditions, and focus fields, without condition-specific parameter tuning, which prevents treatment- or time-dependent segmentation bias.

#### 2.3.2 Segmentation Quality Control

The pipeline incorporates population-level quality control checks designed to detect systematic segmentation failures. Specifically, the pipeline monitors population-level statistics derived from segmentation outputs, including cell size distributions, total cell counts, shape-related statistics, and summary intensity distributions within segmented regions. These statistics are compared across images acquired under the same experimental conditions. Abrupt shifts, strong skewness, or inconsistencies in these distributions are used as indicators of potential segmentation or imaging artifacts.

Based on these statistics, the pipeline identifies common failure modes such as background capture, indicated by excessively large segmented regions; segmentation collapse, characterized by widespread failure to detect valid cellular instances; and imaging-related artifacts, suggested by abnormal intensity distributions within segmented regions. Detection of these failure modes triggers corrective actions within the pipeline and prevents unreliable segmentation results. For failure modes with clear and localized causes such as background capture resulting in excessively large segmented regions, automated parameter adjustments are applied to correct segmentation behavior. While for more complex failure modes, images are flagged for manual inspection and human annotation.

### 2.4 Feature Extraction and Signal Integration Module

The feature extraction and signal integration module transforms segmented cellular instances into quantitative descriptors suitable for longitudinal and condition-dependent analysis. This module operates on the segmentation outputs produced in Section 3.3 and is designed to extract complementary feature types.

In particular, the pipeline extracts two conceptually distinct classes of features: (i) intensity-based features quantify fluorescence signal levels within segmented regions and are particularly sensitive to focus and illumination differences across fields; and (ii) structural features capture cellular organization using morphology and texture descriptors derived from segmentation masks and are generally more robust to illumination fluctuations. Depending on the experimental design, both feature types may be computed from nuclear masks or from derived whole-cell regions, and any fluorescence channel can contribute both features based on assay-specific configuration.

#### 2.4.1 Signal and Feature Extraction

Feature extraction is performed at the single-cell level using lightweight numerical operations and established image analysis libraries, with different feature computations integrated in a modular but unified workflow. Intensity-based features are extracted directly using NumPy operations applied to segmented regions. For each cell, fluorescence intensity statistics (e.g., mean, median, integrated intensity) are computed within the corresponding segmented masks for each relevant channel.

Structural features are computed from the same segmentation masks using functions provided by the *skimage* library. Morphological descriptors, including cell or nuclear area, shape, and eccentricity, are derived from binary masks, while texture features are calculated using gray-level co-occurrence matrix (GLCM)–based statistics computed within segmented regions. These texture metrics capture spatial organization and heterogeneity of signal patterns beyond simple intensity summaries.

All features are computed independently for each segmented cell, producing a cell-level feature table that serves as the fundamental input for downstream aggregation, quality control, and longitudinal analysis.

#### 2.4.2 Feature Type Aware Aggregation

Cell painting images typically include multiple focus fields per imaging channel. Treating each focus field independently can introduce redundancy and amplify noise. To address this, the proposed pipeline performs feature-type–aware aggregation strategies across focus fields. For intensity-based features, aggregation across focus fields is used to reduce sensitivity to focal plane selection and illumination variability. Aggregated values are computed on a per-cell basis using either the mean or median across focus fields. In contrast, for morphological and texture features, a single representative focus plane is selected for each cell based on segmentation confidence, and features are extracted only from that plane. These aggregation strategies are applied uniformly across all experimental conditions and time points, ensuring consistency in longitudinal comparisons.

### 2.5 Downstream summary and visualization

Within SCALE, following feature extraction and aggregation, the resulting per-cell feature tables are summarized to facilitate downstream analysis and interpretation. Summary statistics are computed at the population level for each experimental condition, time point, and feature family. These include measures such as the mean, median, and variability such as variance of features across cells, enabling robust comparisons across conditions while reducing the impact of outliers. Where appropriate, features may be normalized or offset relative to control conditions to facilitate comparative analysis.

Visualization outputs include time-resolved trajectories in reduced feature space, distributions of representative features, and summary heatmaps highlighting changes across conditions. These analyses provide an interpretable link between low-level image-derived features and higher-level phenotypic dynamics, supporting quantitative assessment of temporal responses and experimental perturbations.

### 2.6 Automation and End-to-End Execution

Once the configuration is specified, the pipeline supports a fully automated run from raw images to final analysis tables and figures. Automation ensures that methodological decisions defined in earlier sections are executed deterministically and uniformly across time points and experimental conditions. The framework also supports systematic parameter sweeps and sensitivity analysis by enabling parallel execution under alternative configuration settings. This capability allows evaluation of aggregation strategies, feature subsets, or segmentation backends without modifying core pipeline logic, thereby facilitating robust methodological comparison and reproducible experimentation.

## 3 Pipeline Demonstration on a Chronic Radiation Assay

To evaluate SCALE under a realistic and computationally demanding setting, we applied the pipeline to a longitudinal cell painting dataset generated under chronic low-dose radiation exposure [11]. Radiation was used in this study as a structured and sustained perturbation, with multiple dose levels serving as experimental conditions sampled repeatedly over time. This setting provides a controlled hierarchy of perturbation strength while introducing the temporal variability, imaging heterogeneity, and potential phenotype drift that commonly challenge longitudinal high-content imaging analysis. All segmentation, feature extraction, aggregation, and downstream summarization steps were executed uniformly across dose levels and time points without condition-specific parameter tuning.

### 3.1 Experimental Setting and Dataset Characteristics

Cells were cultured under chronic low-dose radiation exposure at five dose rates (9.4, 4.7, 0.5, 0.05, and 0.01 mGy/hr), with an additional non-irradiated condition serving as the control. Imaging was performed once per week over a nine-week period, yielding a longitudinal dataset that supports dose-stratified temporal comparisons. Each week, multiple 96-well plates were processed using a standardized Cell Painting protocol and imaged with high-content confocal microscopy, ensuring a consistent acquisition workflow across all dose levels and time points.

Four distinct staining configurations were included in this study:

- Canonical Cell Painting panel: the standard six-dye combination (Hoechst 33342, Concanavalin A 488, Phalloidin 568, WGA 555, Mito 641, and 512 nucleic acid stain) capturing nuclear, ER/Golgi, actin, membrane glycoproteins, mitochondria, and RNA-enriched compartments.
- Caspase-modified panel: replacement of the 488 nm Concanavalin A channel with a live-cell caspase-3/7 apoptosis marker, while retaining the remaining canonical channels.
- DNA damage–modified panel (*γ*H2AX only): replacement of the 488 nm Concanavalin A channel with Alexa Fluor 488, a phospho-*γ*H2AX antibody to quantify DNA double-strand break associated signaling.
- Combined apoptosis and DNA damage panel: simultaneous incorporation of caspase-3/7 in the 488 nm channel and phospho-*γ*H2AX in the 647 nm channel, replacing the corresponding canonical markers in both channels.

Representative Week 4 images from the Caspase + *γ*H2AX (647 nm) dual panel across radiation dose levels are shown in Figure 2. The resulting dataset exhibits several characteristics that make it representative of modern longitudinal cell painting studies and computationally non-trivial to analyze:

**Figure 2.**
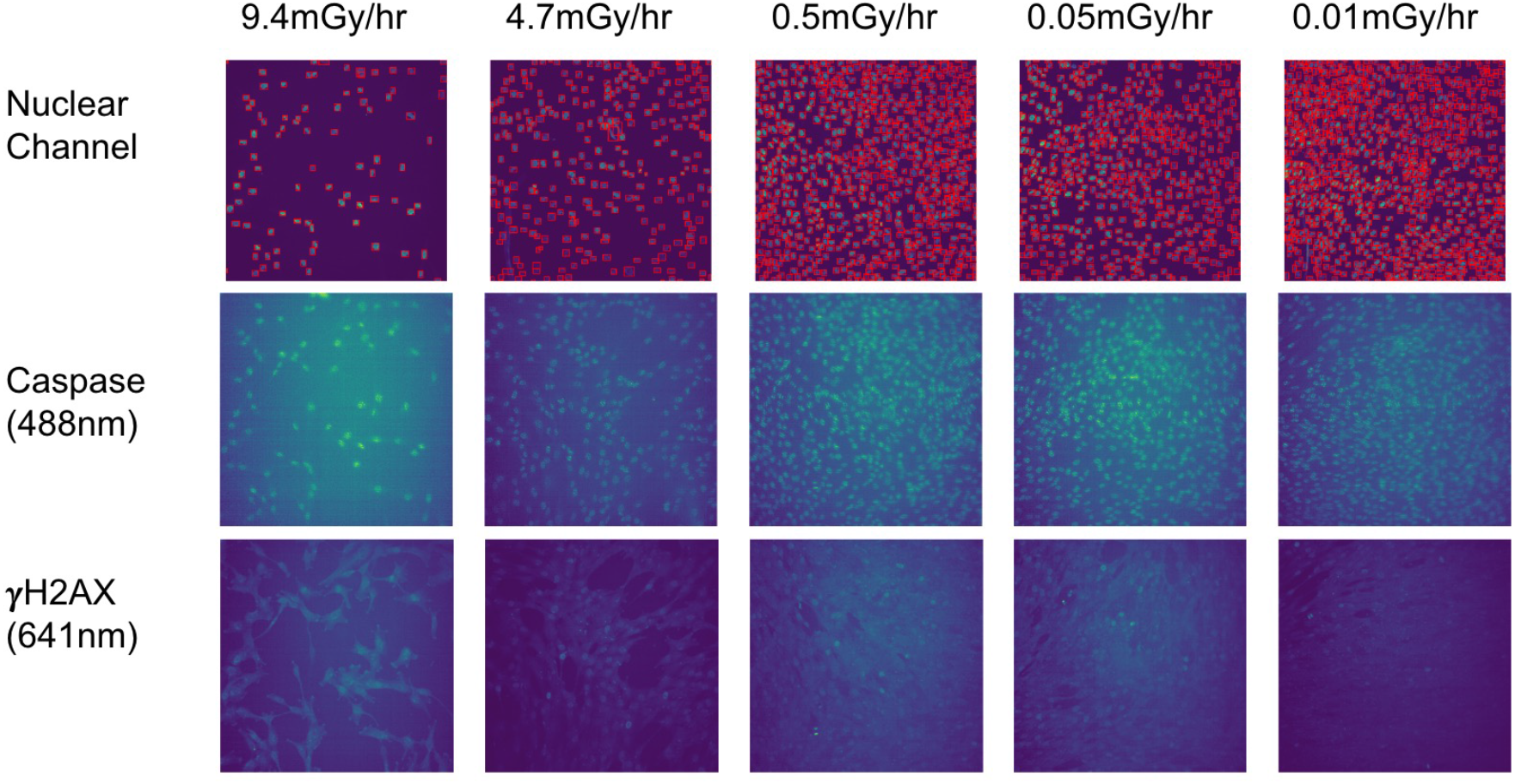
Representative Week 4 images from the chronic radiation Cell Painting assay using the Caspase (488 nm) + *γ*H2AX (647 nm) dual staining panel across five radiation dose rates (9.4, 4.7, 0.5, 0.05, and 0.01 mGy/hr). Columns correspond to dose levels (left to right). Rows correspond to imaging channels: (top) nucleus segmentation masks overlaid on the nuclear channel, (middle) Caspase (488 nm) intensity reflecting apoptosis-associated activity, and (bottom) *γ*H2AX (647 nm) intensity reflecting DNA damage–associated signaling. Visual inspection shows dose-dependent differences in cell density, apoptosis signal intensity, and *γ*H2AX spatial distribution at Week 4. These images illustrate the heterogeneous signal characteristics that motivate stability-oriented, longitudinal analysis.

- Multiple experimental conditions: distinct radiation dose rates treated as categorical perturbation levels.
- Longitudinal sampling: repeated measurements across weeks, requiring consistent processing to enable temporal comparison.
- Multi-channel fluorescence imaging with heterogeneous staining configurations: six primary fluorescent channels were used in the canonical Cell Painting panel to capture diverse cellular compartments, along with brightfield and phase-contrast channels. In parallel wells, modified staining strategies replaced or supplemented selected channels (e.g., alternative markers in the 488 nm and 647 nm channels), resulting in controlled channel heterogeneity across subsets of wells.
- Non-uniform channel semantics across wells: not all wells share identical dye combinations, requiring channel-aware feature extraction and configuration-driven interpretation of feature families.
- Multiple focus fields per well: nine fields per well, introducing across well heterogeneity.
- Z-stack acquisition: fifteen optical sections per field, increasing data volume and necessitating aggregation strategies.
- Large-scale data volume: tens of thousands of images per plate and tens of gigabytes per weekly batch.

Such a dataset places stringent requirements on segmentation stability, feature consistency, focus-field integration, and automation. Manual retuning of analysis parameters across weeks or dose levels would introduce bias and compromise reproducibility. The proposed pipeline is therefore evaluated in this setting to assess its ability to maintain uniform processing and generate comparable longitudinal feature summaries across chronic perturbation conditions.

### 3.2 Configuration Instantiation

#### 3.2.1 Experimental Metadata Encoding

In this study, staining configuration is the top-level analysis identifier: each of the four staining panels (canonical Cell Painting, caspase-modified, *γ*H2AX-modified, and the combined caspase+*γ*H2AX panel) is treated as a distinct pipeline instantiation. Concretely, each staining configuration corresponds to its own channel map, feature-extraction targets, and downstream summaries, and results are not pooled across configurations unless explicitly stated.

For each acquired image, we recorded structured metadata including (i) staining configuration, (ii) channel identity, (iii) focal plane index (z), (iv) field-of-view identifier, (v) well identifier, (vi) radiation dose level, and (vii) week. Dose level and week were encoded as categorical metadata fields, with dose defining the primary experimental grouping variable and week defining the longitudinal axis. Channel and focal-plane labels support channel-specific feature extraction and focus-aware aggregation within each staining-specific pipeline, while well and field identifiers enable hierarchical summaries from cell, field, well to dose × week. The pipeline does not assign intrinsic biological meaning to radiation; dose values are treated generically as condition labels.

#### 3.2.2 Channel and Feature Family Specification

The configuration explicitly defines which channels are used for which feature families under each staining configuration. Across all four staining panels, the 512 nm nucleic-acid channel is designated as the segmentation channel and provides the nucleus-centered masks used throughout the pipeline.

For the canonical Cell Painting panel, the 512 nm channel additionally serves as the primary source for structural features. Specifically, morphology descriptors and GLCM-based texture statistics are computed from the nucleus-centered regions defined by the segmentation masks, yielding structural descriptors that are stable and comparable across weeks and dose levels.

For the modified staining panels, the substituted 488 nm and/or 647 nm channels (e.g., caspase-3/7 and/or *γ*H2AX) are treated as intensity-feature channels. In these configurations, intensity summaries are extracted within the segmentation-derived regions for the modified marker channels. Structural feature extraction is not performed for the modified panels.

#### 3.2.3 Segmentation and Aggregation Parameters

Segmentation was performed using CellSAM as the backend [9]. The primary configurable hyperparameter exposed for this study was the bounding-box confidence threshold, which controls the acceptance of detected nuclear instances. Quality control (QC) for segmentation was implemented using simple, interpretable filters on the resulting instance masks. Specifically, we applied min/max area constraints and an eccentricity filter to identify implausible nuclear masks. Cell-level plausibility filters were applied to each cell. Fields were evaluated using the fraction of instances passing these filters and a minimum valid-cell-ratio criterion. When acceptance criteria were not met, the bounding-box confidence threshold was adjusted and CellSAM segmentation was re-run globally, followed by re-evaluation of QC.

For intensity features, we aggregated signals across the z-stack using a mean-of-means scheme: for each z-plane, we computed the cell-level mean intensity as the average fluorescence within that cell’s segmentation mask on that plane. We then averaged these per-plane cell means across z-planes to obtain a single intensity value per cell. Condition-level mean and variance were subsequently computed by averaging the resulting per-cell intensities across cells, giving each cell equal weight. For the canonical panel, structural features were extracted from a representative focal plane selected based on segmentation confidence. Condition-level summaries likewise include both the mean and variance of each structural feature across cells.

#### 3.2.4 Output and Summary Specification

The primary pipeline outputs are dose × week summary curves for each feature, reporting the mean and standard deviation across cells within each (dose, week) group. These summaries are generated separately for each staining configuration and serve as the main analysis-ready representation. In addition to the raw summaries, we also generate control-normalized trajectories for longitudinal comparison. For each week and staining configuration, feature summaries at each dose are expressed relative to the corresponding non-irradiated control, yielding dose-dependent change curves anchored to a common reference.

### 3.3 Feature-Type–Aware Longitudinal Behavior

We summarize what the pipeline reveals when feature families are tracked longitudinally under chronic radiation. The key observation is a feature-type–dependent shift in dose information over time: early in the exposure course, dose stratification is expressed most strongly in intensity-based marker readouts, whereas at later weeks, dose-dependent separation is better preserved by texture-derived structural descriptors from the canonical panel.

#### 3.3.1 Intensity-based Features

For intensity-based readouts, the pipeline produces dose × week trajectories for the marker channels in each modified staining configuration and reports both the raw mean and std across cells and control-normalized counterparts (percentage difference from the non-irradiated control at the same week). Figure 3 summarizes the *control-normalized* trajectories for four marker settings: *γ*H2AX(488)-only, Caspase(488)-only, Caspase(488) and *γ*H2AX(647) in the dual panel.

**Figure 3.**
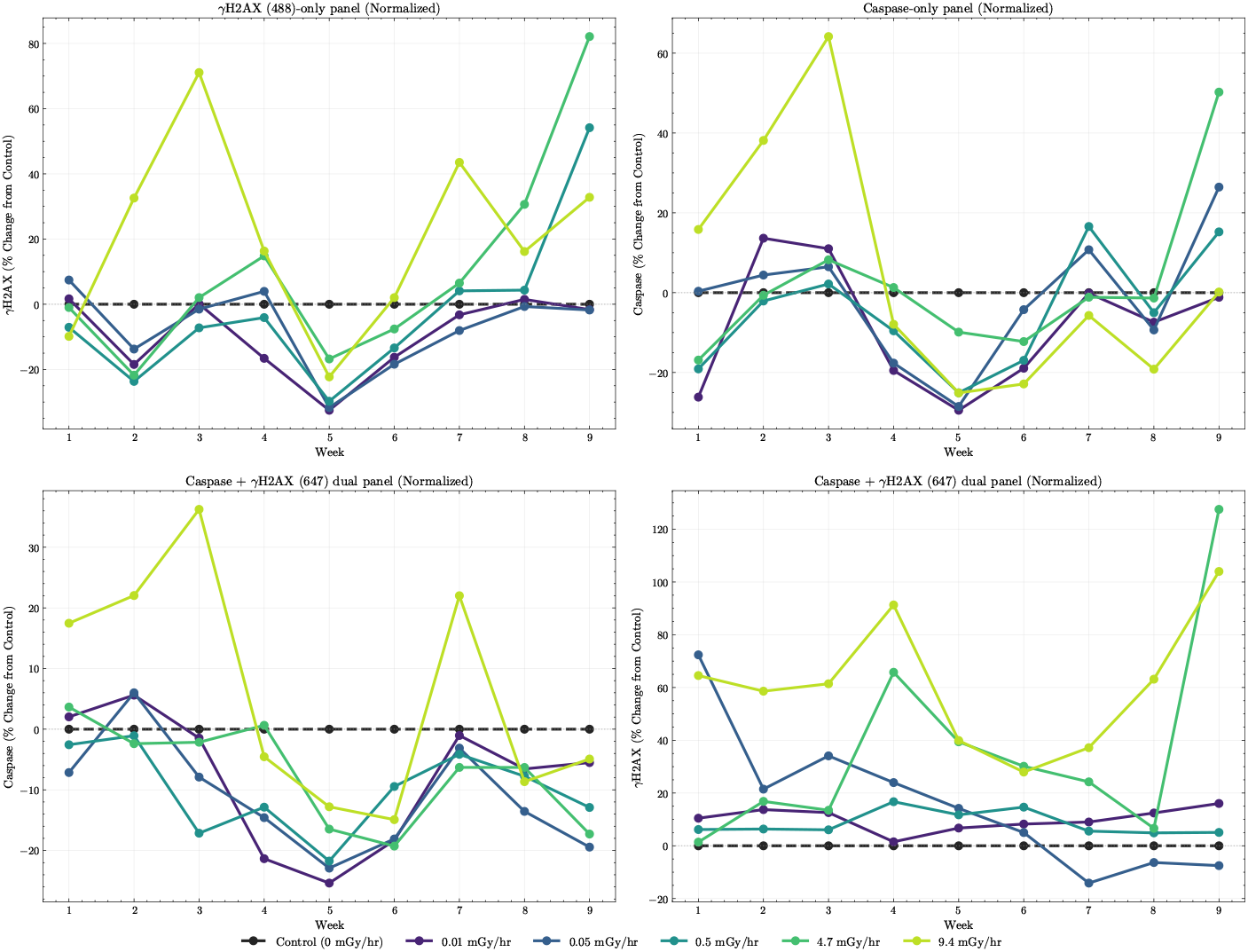
Control-normalized intensity trajectories for modified marker panels. Dose × week change curves (percent change from the non-irradiated control at the same week) for four intensity-marker settings: (top-left) *γ*H2AX(488)-only panel, (top-right) Caspase(488)-only panel, (bottom-left) Caspase(488) in the Caspase+*γ*H2AX(647) dual panel, and (bottom-right) *γ*H2AX(647) in the dual panel. Curves are shown for control and five dose rates (0.01, 0.05, 0.5, 4.7, 9.4 mGy/hr). Each point represents the weekly mean across cells after z-stack aggregation at the cell level; trajectories illustrate strong early dose-dependent separation for intensity markers and partial convergence in mid-to-late weeks for several channels, with marker-specific differences in persistence and dynamic range.

Across panels, intensity differences are most pronounced during the early weeks of exposure and generally scale with dose, consistent with a strong early dose-dependent response. In Fig. 3, higher-dose conditions (4.7 and 9.4 mGy/hr) show clearer separation from lower-dose groups during Weeks 1–3 in both the *γ*H2AX-only and Caspase-only settings, while many trajectories exhibit reduced separation or partial convergence during mid-to-late weeks (notably around Weeks 5–8) despite continued exposure. The dual-panel readouts further highlight that dose-dependent trends can differ across markers: Caspase trajectories show comparatively smaller dynamic range than *γ*H2AX(647), whereas *γ*H2AX(647) displays sustained positive deviations at higher doses and a pronounced late-week increase in the highest-dose conditions.

#### 3.3.2 Texture and Structural Features

In contrast to intensity markers, structural features exhibit more persistent dose-dependent separation over the nine-week exposure course, as shown in Fig. 4. The normalized dose × week trajectories reveal that morphology descriptors (e.g., area, perimeter, eccentricity, solidity) and texture metrics (e.g., GLCM correlation, GLCM energy, and GLCM homogeneity) retain distinguishable dose stratification beyond the early response phase. Notably, several texture descriptors, most prominently GLCM energy and GLCM homogeneity, show sustained separation between higher-dose (4.7 and 9.4 mGy/hr) and lower-dose conditions during mid-to-late weeks, even when intensity-based trajectories converge. While early weeks still display dose-dependent shifts in both morphology and texture, structural features maintain clearer relative ordering across dose levels in Weeks 6–9. In particular, GLCM-based features exhibit systematic negative shifts for higher doses at later weeks, whereas lower-dose groups cluster closer to control, indicating persistent alterations in spatial organization rather than transient intensity amplification.

**Figure 4.**
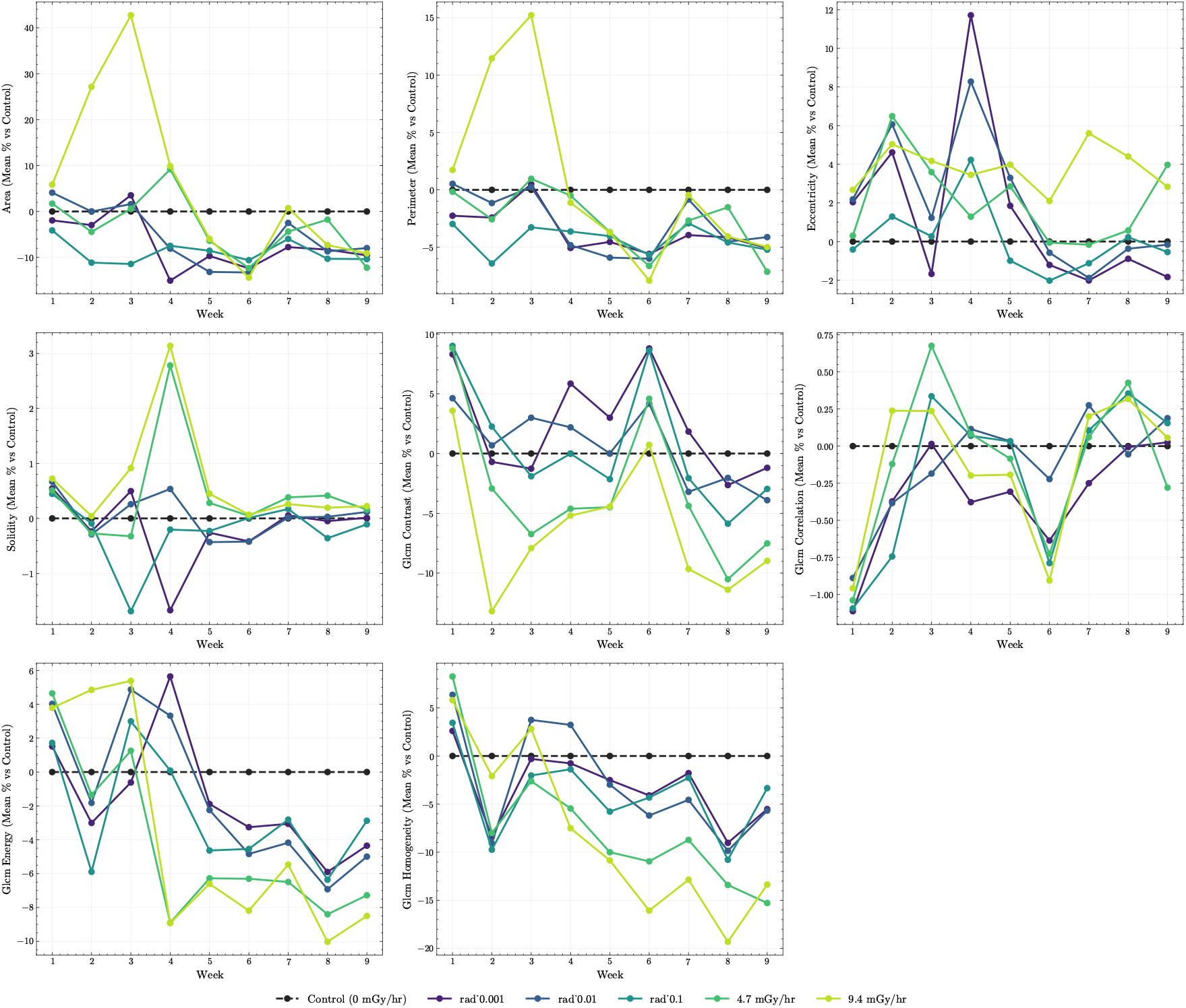
Normalized structural feature trajectories (dose × week). Control-normalized mean trajectories for representative morphology and texture descriptors from the canonical Cell Painting panel, including area, perimeter, eccentricity, solidity, GLCM contrast, GLCM correlation, GLCM energy, and GLCM homogeneity. Curves are shown for control and five dose rates (0.01, 0.05, 0.5, 4.7, 9.4 mGy/hr). Several texture metrics—particularly GLCM energy and homogeneity—maintain clearer dose-dependent separation during mid-to-late weeks compared to intensity-based features, indicating persistent alterations in nuclear organization under chronic exposure.

From a pipeline perspective, these results demonstrate that texture-based descriptors provide a more stable longitudinal readout of cellular organization under chronic perturbation. Unlike intensity features, which are more sensitive to focal-plane selection and illumination variability, structural features anchored to nucleus-centered masks capture organizational changes that remain separable across time. This supports the feature-type–aware design of the pipeline, in which intensity and structural features are processed and interpreted as complementary longitudinal signals.

## 4 Conclusion and Discussion

In this work, we introduced SCALE (Stable Cell Painting Analysis for Longitudinal Experiments), a stability-oriented, configuration-driven framework to longitudinal Cell Painting analysis by integrating segmentation, automated QC, feature extraction, aggregation, and standardized reporting within a single configurable workflow. Using a nine-week chronic radiation assay as a test case, the pipeline executes uniformly across dose levels and time points to produce analysis-ready dose × week trajectories, and the resulting summaries illustrate a feature-type–dependent longitudinal behavior in which intensity-based markers show stronger early dose stratification while nucleus-centered texture descriptors (e.g., GLCM energy and homogeneity) preserve dose-dependent separation more consistently at later weeks.

Longitudinal Cell Painting is fragile when focal-plane selection, ad hoc filtering, or inconsistent aggregation drift over weeks; the proposed pipeline mitigates these risks by making experiment structure explicit in configuration and enforcing condition-agnostic processing rules and standardized outputs. The current implementation can be strengthened in several ways: (i) extend QC beyond segmentation plausibility to include feature-level stability checks that monitor distribution shifts across weeks, plates, and channels; (ii) add configuration-driven sensitivity analyses for aggregation/normalization choices and report richer records (QC/provenance summaries and optional cell-level tables) to improve debugging, reproducibility, and Interpretability.

## Appendix

To complement the control-normalized trajectories presented in the main text, we provide the corresponding *raw* dose × week summaries. These supplementary figures serve as audit artifacts that expose absolute feature magnitudes and within-condition variability over time, ensuring full transparency of the aggregation procedure. For all features, cell-level measurements were first computed under a fixed SCALE segmentation and aggregation scheme (including z-stack integration for intensity features), and then summarized across cells within each (dose, week) group. No week-specific or condition-specific parameter adjustments were introduced during processing.

**Figure S1.**
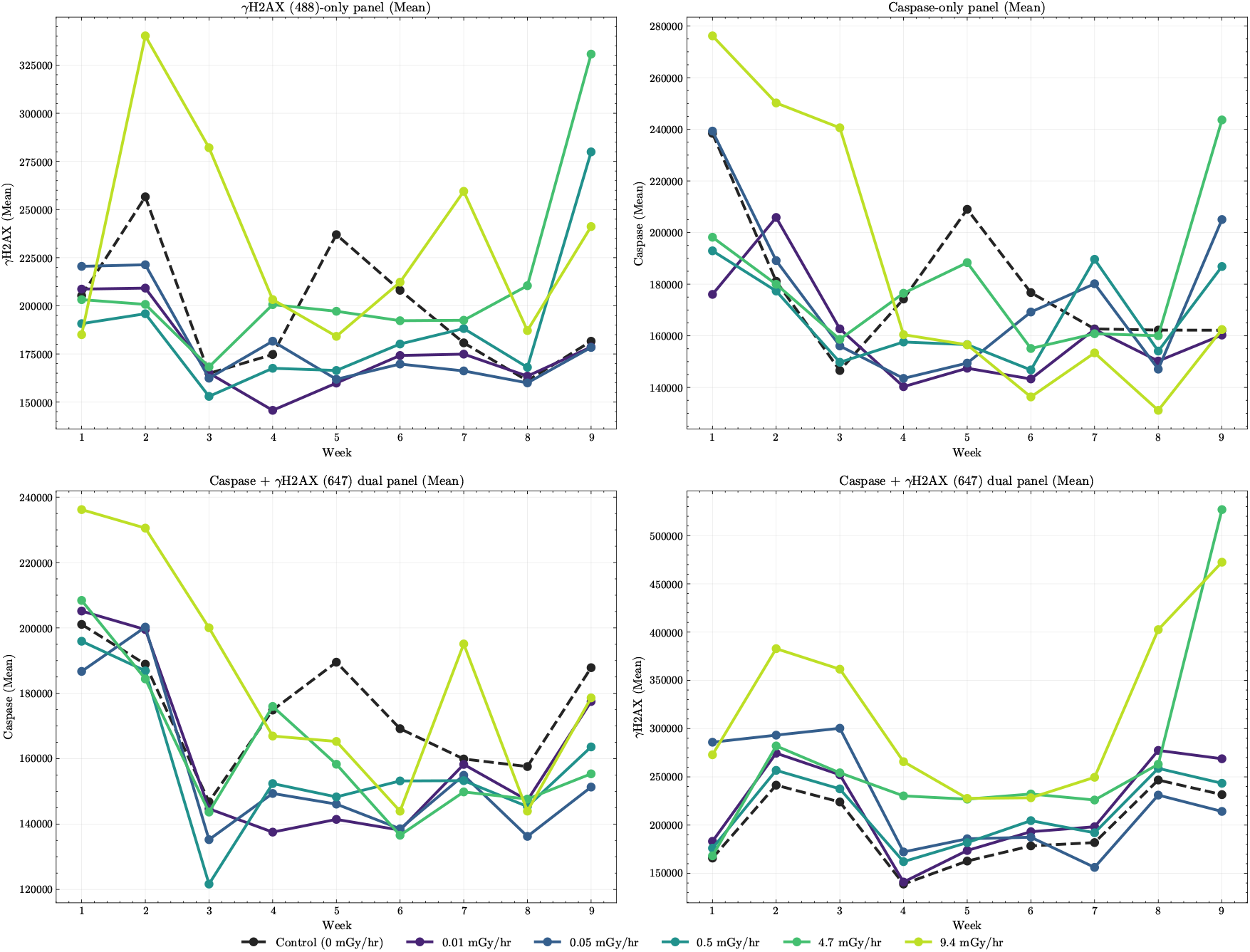
Raw mean intensity trajectories (dose × week). Weekly dose × week trajectories of the *mean* cell-level intensity (a.u.) for four intensity-marker settings: (top-left) *γ*H2AX(488)-only panel, (top-right) Caspase(488)-only panel, (bottom-left) Caspase(488) in the Caspase+*γ*H2AX(647) dual panel, and (bottom-right) *γ*H2AX(647) in the dual panel. Curves are shown for control and five dose rates (0.01, 0.05, 0.5, 4.7, 9.4 mGy/hr).

**Figure S2.**
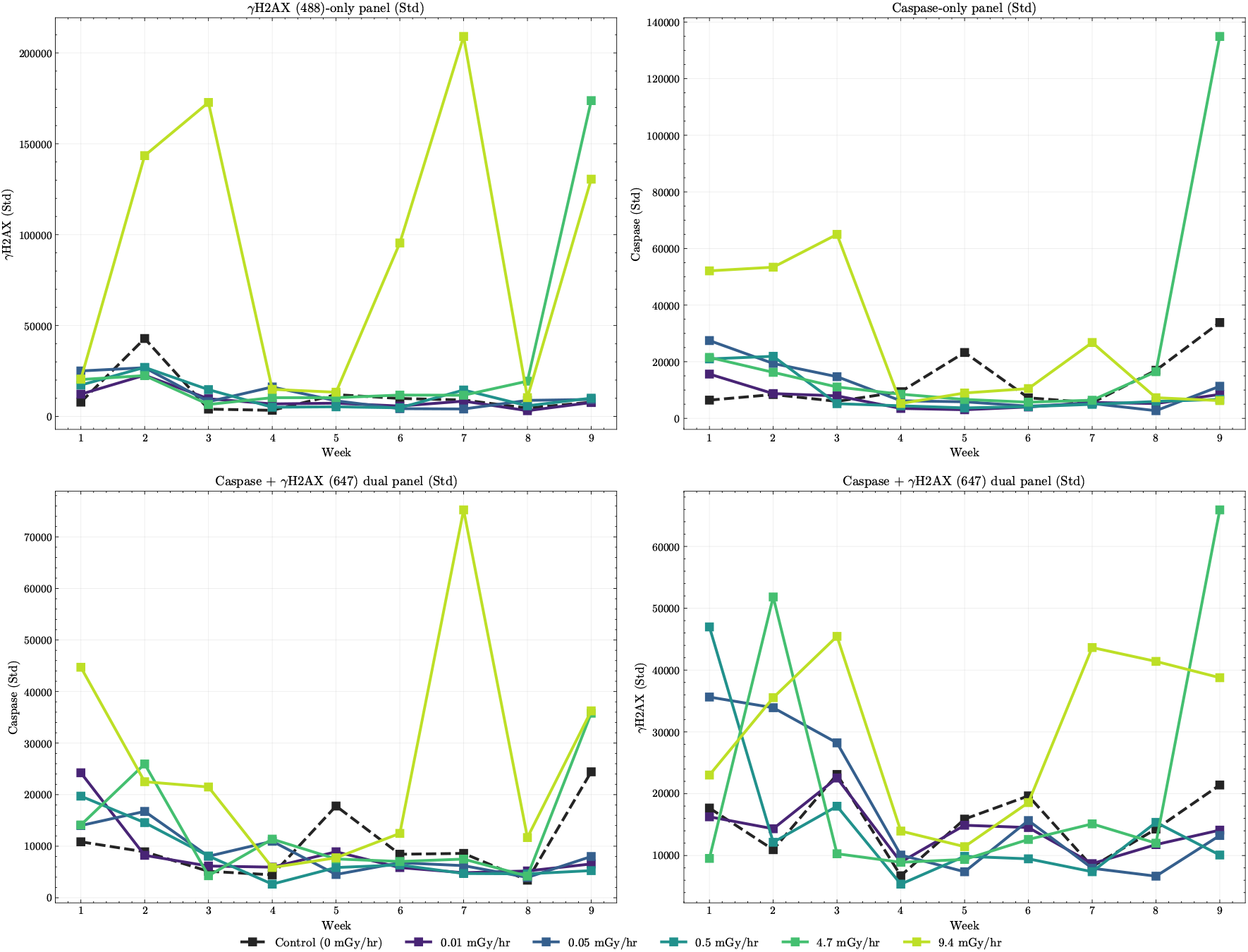
Raw intensity dispersion trajectories (dose × week). Weekly dose × week trajectories of the *standard deviation* of cell-level intensity (arbitrary units) for the same four intensity-marker settings as Figure S1. Standard deviation is computed across cells from per-cell z-aggregated intensities (mean-of-means across z), and provides a compact summary of within-condition heterogeneity that complements the mean trajectories.

**Figure S3.**
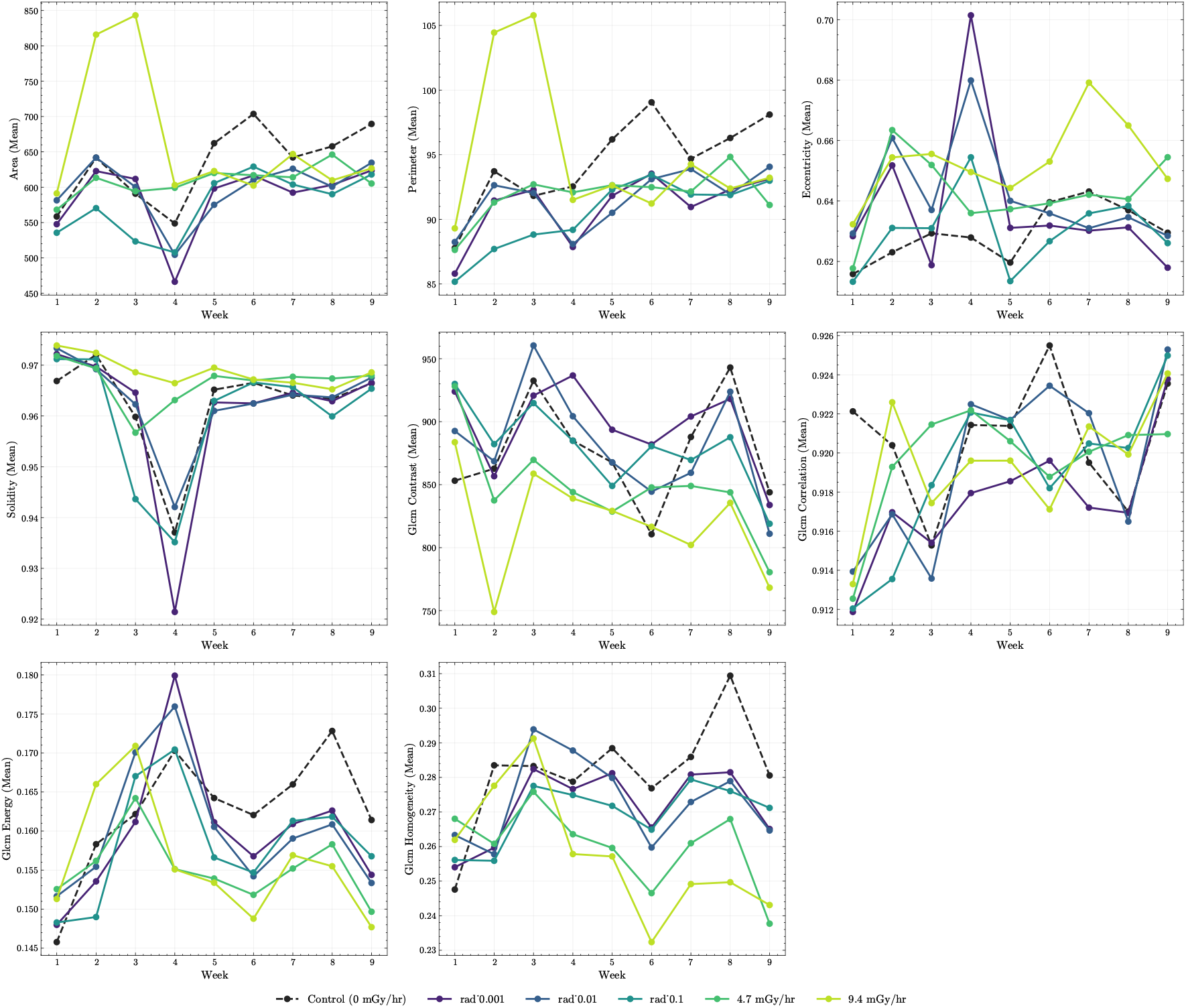
Raw mean structural feature trajectories (dose × week). Weekly dose × week trajectories of the *mean* structural features (arbitrary units) derived from the canonical Cell Painting panel, including morphology descriptors (area, perimeter, eccentricity, solidity), and GLCM texture metrics (contrast, correlation, energy, homogeneity). Each point represents the mean across cells of per-cell aggregated measurements under a fixed segmentation and aggregation scheme. Curves are shown for control and five dose rates (0.01, 0.05, 0.5, 4.7, 9.4 mGy/hr).

**Figure S4.**
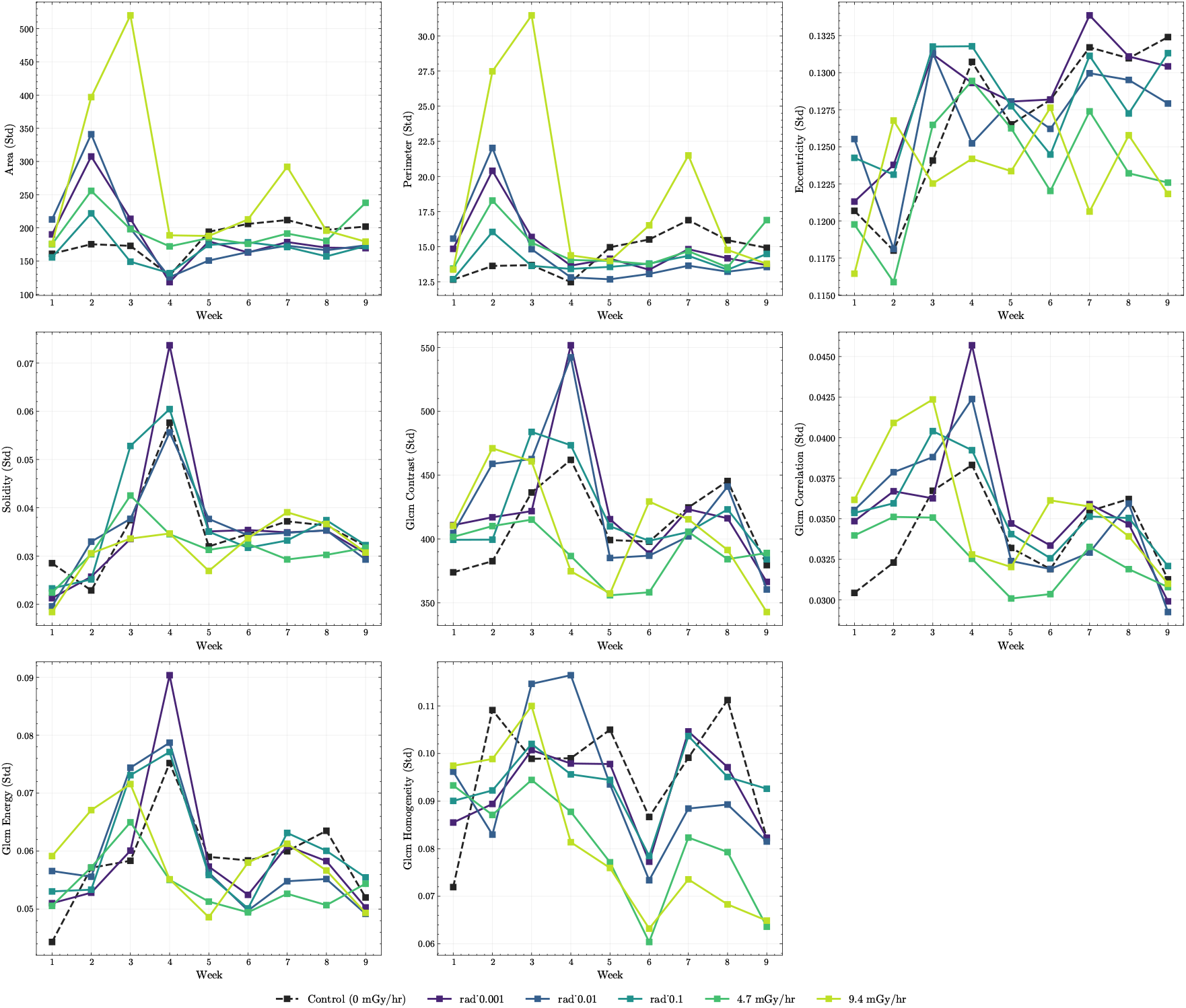
Raw dispersion of structural features (dose × week). Weekly dose × week trajectories of the *standard deviation* across cells for the same structural features shown in Figure S3. Standard deviation summarizes within-condition heterogeneity and complements the mean trajectories, enabling assessment of dose-dependent variability in morphology and texture over time.

